## Supplementary Information for "Paninvasion severity assessment of a U.S. grape pest to disrupt the global wine market"

**This PDF file includes:**

Supplementary Methods
Supplementary Table 1
Supplementary References

**Supplementary Methods**

Below, we provide additional details for terminology and methods for the analyses conducted in our study.

*Term Definitions*

- **alignment correlation**—multivariate relationship among invasion potentials.
- **establishment potential**—likelihood of a region to contain suitable habitat for transported individuals of a non-native species to form a spreading population.
- **impact potential**—likelihood of a region to experience negative economic effects from an established non-native species.
- **invasion potentials**—likelihoods of a species to move through stages in an invasion process across regions[1]. We focus on the main stages: transport, establishment, impact.
- **MaxEnt**—abbreviation for maximum entropy, a presence-only SDM methodology that uses machine-learning to estimate the probability distribution of maximum entropy based on environmental variables and species occurrence records[2,3].
- **paninvasion**—invasion of a species at the global scale that disturbs a global economic market.
- **paninvasion risk**—the likelihood of a regional invasive species to become a globally invasive species and cause economic repercussions.
- **paninvasive species**—globally invasive species that goes through the three main invasion stages and thus can disturb global economic markets.
- **phylloxera**—*Daktulosphaira vitifoliae* is grapevine root pest native to North America that was responsible for the Great Wine Blight of the late 1800's, which was the largest economic disturbance to the global wine market ever recorded. The disruption was mitigated by widespread planting of European vines that were grafted to North American grapevine root stocks. The paninvasion of phylloxera continues to this day[4,5].
- **species distribution model (SDM)**—spatial model used to predict the environmental niche, habitat suitability, and establishment potential of a species.
- **spotted lanternfly (SLF)**—*Lycorma delicatula* is a planthopper native to China, Vietnam, and India. It invaded South Korea and Japan in the early 2000's and the

northeastern U.S. ca. 2014. It is known to feed on >100 different host species, including grapes[6,7].

- **transport potential**—likelihood of a region to have an introduction of a non-native species.
- **tree of heaven (TOH)**—*Ailanthus altissima* is a paninvasive deciduous tree that is native to China, Taiwan, and northern Korea, but has been spread globally. It is a highly preferred host for SLF and may determine SLF establishment potential.

*Supplementary methods: Confirmation of relationship between import tonnage and SLF invasion status for transport potential*

The prevailing hypothesis on SLF transport potential is that regions that import more tonnage of commodities from the invaded U.S. region also import more total tonnage of goods and trade infrastructure (e.g., cargo containers, pallets, railcars) that inadvertently transport SLF egg masses long-distances. SLF propagules have been found hitchhiking on and in shipments of pharmaceutical containers, baking ingredients, paint shipments, building materials, boxes of pumpkins, pallets and many other commodities[6,8–12]. To test if total tonnage can explain the current spread of SLF, we fit two logistic regressions with our metric of transport potential based on total tonnage as the covariate. This metric was the $\log_{10}$ of the average annual metric total tonnage imported between 2012 and 2017 from U.S. states invaded by SLF (main text Fig. 3). We regressed the presence/absence of established populations and regulatory incidents (i.e., has a state experienced and reported any observations of SLF, dead, moribund, or alive, independent of the presence of established populations?). For both establishment and regulatory incidents, the relationship between SLF-status and our measure of transport potential was significant, thereby providing support for our estimate of SLF transport potential (Supplemental Table 1). These results suggest that total tonnage of imports is a suitable proxy for transport potential until new metrics are developed that include refined pathway analyses.

*Supplementary methods: Modeling establishment potential and the influence of chilling periods for diapause*

We estimated establishment potential as an ensemble from three global species distribution models (SDMs): a multivariate SDM of TOH (*sdm_toh*), a multivariate SDM of SLF (*sdm_slf1*), and a univariate SDM of SLF that modeled SLF presence on the predicted values from *sdm_toh (sdm_slf2)*. Models were constructed with MaxEnt ver. 3.4.1 by following best practices for estimating unbiased niche models[3,13,14]. We first queried GBIF for TOH and SLF presences on October 20, 2020. For TOH, 67,100 records were obtained and for SLF 3,180 records were obtained[15]. Records were checked for errors, duplicate records removed, and the remaining records were rarefied (spatially filtered) by omitting records <10 km from each other to reduce bias from spatial autocorrelation[16,17]. The result was 8,022 unique, error checked TOH presence records and 325 unique, error checked SLF presence records. Thus, *sdm_toh* was built on 8,022 TOH global presence records, and *sdm_slf1* and *sdm_slf2* were built on 325 SLF presence records (see our research compendium and Dryad repository for the data, https://github.com/ieco-lab/slfrsk and https://doi.org/10.5061/dryad.msbcc2g1b).

To find the best models that explained TOH and SLF presences, we started with 22 potential covariates hypothesized to influence SLF and TOH global distributions. The covariates included 20 topographic and bioclimatic variables from WorldClim, which is a standard database of covariates used in global SDMs[18,19]. WorldClim has also been used in two previous SDMs for SLF[20,21]. In addition to these 20 covariates, we added Global Forest Canopy Height[22] because SLF feeds on multiple tree species[23], and Global Access to Cities[24] because TOH and SLF are often established along transportation networks[9]. We analyzed these covariates to identify an uncorrelated subset to include in final best-fit SDMs with low model collinearity. To do this, we calculated pairwise Pearson correlations among the 22 covariates, and fit each covariate to SLF and TOH in univariate SDMs (i.e., 44 models in total). We then compared covariates that were highly correlated and retained only the covariates that fit best to the TOH and SLF presences. This reduction of potential covariates resulted six minimally correlated covariates (pairwise absolute Pearson correlations <0.70) that we fit in our models: annual mean temperature (BIO01), mean diurnal temperature range (BIO02), annual precipitation (BIO12), precipitation seasonality (BIO15), elevation (ELEV), and access to cities (ATC).

We fit *sdm_toh* and *sdm_slf1* with these six covariates; *sdm_slf2* was fit from the *sdm_toh* predicted values. The three models were fit under default settings of the MaxEnt program except for the following changes: (1) all features were enabled but still set to "Auto Features", (2) response curves were created, (3) variable importance was measured via jackknifing (we did not do this for *sdm_slf2* because it was a univariate model), (4) the threshold rule was set to “Minimum Training Presence”, and (5) the number of replicates was set to five for SLF and ten for TOH. This last modification sets the number of *k*-fold cross-validation replicates and determines the test proportion from *k*, thus we validated the three models with *k*-fold cross-validation via evaluation of the receiver operating characteristic of the AUC (area under the curve) and omission error[2,25–27]. For AUC, the fraction of true positives relative to type I error (positive background points) is compared at all possible thresholds for each model[2,25]. The resultant AUCs were assessed relative to a random model where AUC = 0.50, such that values close to 1.00 indicate strong model performance and those ≤ 0.50 suggest poor performance[25]. Given presence only data, measured AUC cannot reach 1.00, but model AUCs that approach 1.00 are considered to perform well[2,28]. Given concerns with model evaluation with AUC[29–31], we also confirmed model performance with average omission error, which is the proportion of presence point(s) predicted with suitability less than the threshold averaged across replicates[26,27].

All three models performed well according to AUC and omission error. Models yielded test AUC values >0.75 while boasting average test omission error rates <0.01, indicating that each model performed better than random and identified areas of known species presence as suitable for the cross-validation partitions. *sdm_toh* had a slightly lower AUC (0.7779) and omission error (0.0003) than *sdm_slf1* (AUC = 0.9828, omission = 0.0064) and *sdm_slf2* (AUC = 0.9675, omission = 0.0032). For both multivariate SDMs, we compared the variable contributions for congruence. The top four contributing variables were the same for both models (ATC, BIO01, BIO12, and BIO15 in descending order). The remaining two variables (ELEV and BIO02) contributed <2% in each model, with ELEV contributing more in *sdm_toh* and BIO02 contributing more in *sdm_slf1*. For *sdm_toh*, two other variables, BIO12 and BIO15 also contributed <2% each but still contributed more than ELEV and BIO02 (for a more detailed comparison, see our research compendium, https://ieco-lab.github.io/slfrsk/).

We averaged our three best-fit models to produce one ensemble image at the 30 arcsecond resolution, and intersected this image with state and country polygons[13]. We then calculated summary statistics for the ensemble pixels within each state and country (mean, median, and maximum). The R function we wrote to perform this task, extract_enm2(), is available with the R companion package, slfrsk (see https://github.com/ieco-lab/slfrsk). Establishment potential for the 50 U.S. states and 223 countries was estimated as the maximum pixel value for each state and country. Results and conclusions with mean and median pixel values instead of max were qualitatively similar (see https://ieco-lab.github.io/slfrsk/).

Although our work suggests widespread establishment potential, SDM-based establishment potential might overestimate suitability in warmer climates if SLF require a chilling period to initiate diapause to complete development[32]. However, recent work suggests that while SLF can diapause as eggs in the invaded U.S. region, native populations across China include sub-tropical regions that do not provide the colder temperatures necessary for completing diapause[33], and SLF in the U.S. do not require diapause to develop[34]. Indeed, under lab conditions, eggs in the U.S. that do not undergo diapause exhibit higher survivorship than those that do undergo diapause[35]. This observation suggests that our global ensemble model does not overestimate SLF establishment potential and instead may be a conservative estimate, especially for warmer regions (main text Fig. 4).

In summary, our estimate of SLF global establishment potential was based on an ensemble of models for SLF and TOH environmental suitabilities. Two previous estimates of SLF global establishment potential have been published but did not include TOH, were not ensemble estimates, and were not built on as many presence records[20,21]. These other estimates also did not include an anthropogenic covariate like Global Access to Cities[24], which we found to be important in determining TOH and SLF environmental suitability. Finally, although our estimate of SLF establishment potential is broadly like these previous estimates (as observed by comparing our map to theirs), it differs in three key ways: we provide our estimate in a finer resolution, our estimate differs across globally important viticultural regions, and we provide the data as open access. To visualize and download our estimate please see our Google Earth Engine app (https://ieco.users.earthengine.app/view/ieco-slf-riskmap).

**Supplementary Table 1** Logistic regression of spotted lanternfly (SLF) status on trade with established U.S. states as average annual metric total tonnage demonstrates a significant relationship for all U.S. states and Washington D.C. Trade with established states predicts both presence or absence of established SLF populations and record of SLF regulatory incidents (identification of SLF, deceased, moribund, or alive). Logistic regression model coefficients are shown above with standard error below in parentheses.

| | Establishment Status | Regulatory Status |
|---|---|---|
| $Log_{10}$(average annual metric tonnage) | 5.64$^{***}$ | 3.10$^{***}$ |
| | (2.03) | (0.90) |
| Constant | -42.74$^{***}$ | -22.64$^{***}$ |
| | (15.27) | (6.49) |
| *Observations* | 51 | 51 |
| *Log likelihood* | -9.80 | -18.15 |
| *Akaike information criterion* | 23.61 | 40.30 |
| *Notes:* | $^{***}P < .01$ | |
